# Shared computational principles for mouse superior colliculus and primate population orientation selectivity

**DOI:** 10.1101/2025.02.22.639626

**Authors:** Austin Kuo, Justin L Gardner, Elisha P Merriam

## Abstract

While the mouse visual system is known to differ substantially from the primate, if the two systems share computational principles, then generalization of results across species may still be possible. One prominent difference is that orientation selectivity is found in mouse superficial superior colliculus (SC), but is not commonly observed in primate SC. Nevertheless, there may be conservation of computational principles if orientation selectivity in mouse superficial SC displays similar properties to primate primary visual cortex (V1), such as invariance to differences in other stimulus dimensions. However, a recent calcium (Ca^2+^) imaging study revealed a population map for stimulus orientation in mouse superficial SC that changed with stimulus properties such as size, shape and spatial frequency, in apparent contradistinction to computational principles for orientation selectivity in primates. To reconcile mouse and primate mechanisms for orientation selectivity, we constructed computational models of mouse superficial SC populations with fixed, stimulus-invariant receptive fields (RFs) classically used to describe neural RFs in monkey lateral geniculate nucleus (LGN) and V1. At preferred spatial frequencies, model RFs exhibited stronger responses where the aperture and gratings were differently oriented, while at non-preferred frequencies, orientation selectivity reversed, matching the imaging data. We provide an intuitive explanation by visualizing stimulus-RF interactions in the spatial frequency domain. Intrinsically oriented RFs were unnecessary to explain much of the imaging data, but modeling of single units suggests a possible subpopulation of intrinsically orientation-selective cells. In summary, our population modeling approach provides a parsimonious explanation for stimulus-dependent orientation selectivity consistent with well-established results from sensory neurophysiology. More broadly, we provide a population modeling framework for establishing shared computations across species.

## Introduction

A foundational finding of sensory neurophysiology is that selectivity for basic stimulus properties is invariant to different stimulus manipulations and behavioral states. A prominent example of this is orientation selectivity in V1. Orientation selectivity arises from feedforward anatomical projections from the LGN to visual cortex^1–4^. The resulting neural selectivity is invariant to changes in stimulus contrast^5,6^, presence of suppressive stimuli^7^, and attentional state^8^. This extensive line of research suggests that selectivity can be considered a fixed property that does not change with stimulus context.

A recent study^9^ found orientation selectivity depended on the visual stimulus. Ca^2+^ imaging measurements from mouse superficial superior colliculus (sSC) neurons demonstrated maps of orientation-selective responses localized at retinotopic locations corresponding to stimulus edges. Furthermore, orientation-selective responses at the same locations along the stimulus edge shifted to show preference for orthogonal orientations with increasing stimulus spatial frequency. These changes in selectivity with stimulus properties might suggest that the computational principles underlying RF properties known from primate research are not shared by mouse SC.

Here we ask whether the changing orientation selectivity reported in mouse sSC can be reconciled with fixed receptive field (RF) properties. We simulated neural responses through a computational model to determine whether the prior imaging results^9^ necessarily imply that mouse sSC neurons have RFs that change their intrinsic orientation selectivity. We constructed a simple spatial stimulus energy model of mouse sSC using RFs with fixed properties based on previously reported measurements^10–15^. We found that our model built entirely of fixed RFs was sufficient to reproduce the phenomenon of changing orientation selectivity. Moreover, we found that center-surround RFs were sufficient to produce orientation maps, suggesting that orientation selectivity measured in a neural population can be an emergent property from inherently non-oriented single-unit RFs. However, our modeling framework does not preclude the existence of RFs with inherently oriented structure in mouse sSC, as we found that simulated V1-like RFs could also produce changing population orientation preference maps and were needed to explain a small population of preference-maintaining neurons from the imaging data. As described below, many of the changes in orientation selectivity with spatial frequency are readily explained by considerations of both stimuli and RFs in the spatial frequency domain. These results highlight that spatial frequency selectivity is essential for producing the phenomenon of changing orientation selectivity at the population scale.

## Results

### Center-surround RFs with fixed properties produce radial orientation preferences

To determine what aspects of the imaging results can be accounted for by RFs not intrinsically selective for stimulus orientation, we built a computational model using only circular-symmetric, center-surround RFs. RFs tiled the visual field with size (**σ**_center_: 1.5°, **σ**_surround_: 11.4°) and peak spatial frequency preference (0.04 cycles/°) aligned with previously reported measures of RF parameters in mouse sSC^11–15^. Stimuli were oriented gratings with parameters for orientation, spatial frequency, aperture shape and size matching those used in prior imaging experiments^9^ (Fig. 1b). Model single-unit RF responses were obtained by computing the dot product between the stimuli and RFs, followed by thresholding and summing resulting values across pairs of On-center, Off-surround and Off-center, On-surround RFs (Fig. 1c).

**Figure 1.**
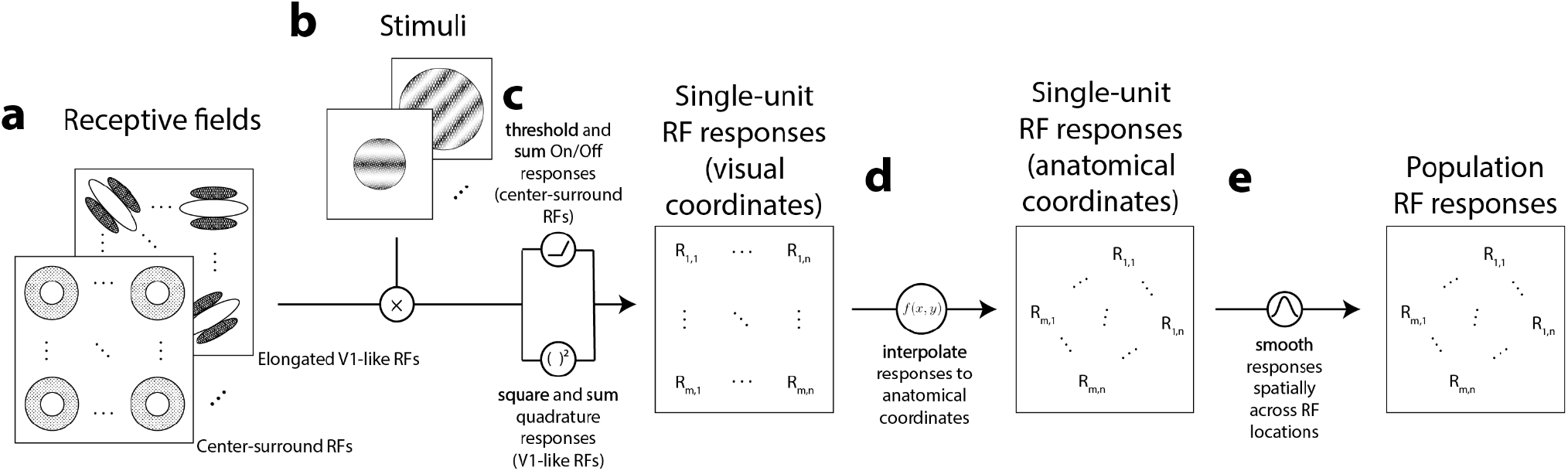
Model of single-unit and population responses from simulated RFs. **A)** Center-surround and randomly oriented V1-like RFs tiled a dense grid of locations; the grid of RF centers extended beyond the extent of all stimuli. **B)** Stimuli were sinusoidal gratings with varying orientations, phases, and spatial frequencies; specific parameters depended on simulation. Stimulus apertures varying in size and shape matched those used previously^9^. **C)** Stimuli and RFs were pointwise multiplied for each pairwise combination of stimulus and RF parameters and combined across RF On-Off pairs or RF phase to generate single-unit RF responses. **D)** RF responses for evenly sampled locations in anatomical space were estimated from grid of RF responses in visual coordinates using natural neighbor interpolation. **E)** Single-unit RF responses in anatomical space were spatially smoothed using a circular symmetric Gaussian kernel, resulting in a set of simulated population RF responses.

To compare the results of our model to wide-field Ca^2+^ imaging data, we spatially transformed the simulated responses to better match the anatomical coordinates in mouse sSC (Fig. 1d). Using maps of visual and anatomical space from prior imaging data^9^ (Fig. 2a), we constructed a set of functions that estimated visual coordinate locations from anatomical coordinate locations (Fig. 2b). This transformation resulted in a roughly 90° clockwise rotation and spatial warping from visual to anatomical space (Fig. 2a, inset transformation, right). Single-unit RF responses at a simulated grid of anatomical coordinates were then estimated by transforming RF responses for corresponding coordinates in visual space (Fig. 2c). Each pixel in the Ca^2+^ imaging measurement reflected the joint activity of a population of neurons. Hence, to simulate these measurements, we locally averaged responses from our simulated RFs in anatomical coordinates using a Gaussian filter with a standard deviation of 0.44° (~0.0083 mm).

**Figure 2.**
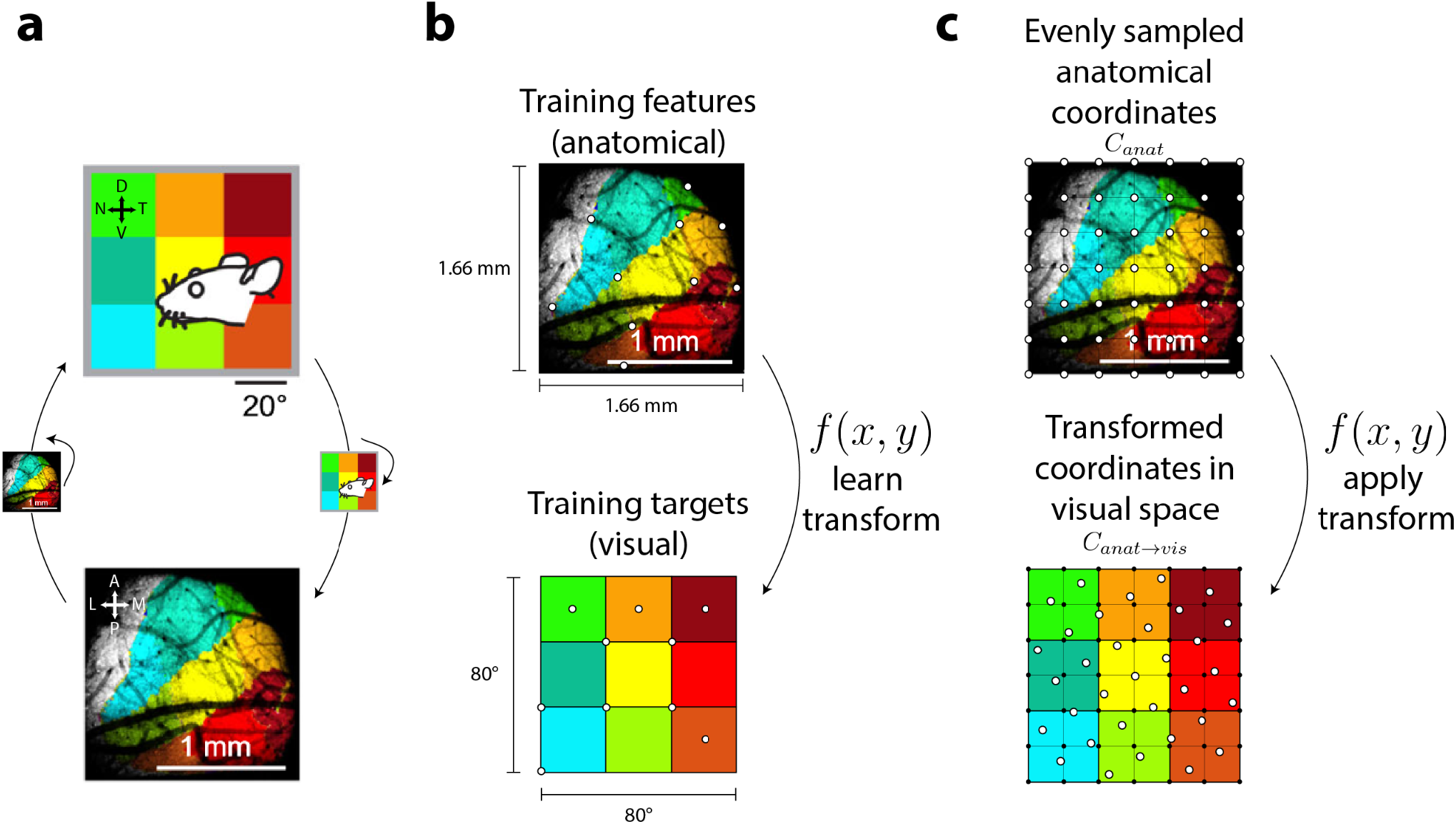
Aligning anatomical and visual coordinates. **A)** Map of visual space (top) and corresponding aligned coordinates in anatomical space for an example mouse (adapted from Fig. 1, Ref. 9). **B)** Anatomical space mapped onto visual space using natural neighbor interpolation to learn transforms from anatomical input coordinates to visual output coordinates. Functions approximating these transforms were learned using a set of training features (top, white points) and a set of training targets (bottom, white points). **C)** Once learned, functional transforms were applied to convert given points in anatomical space (top, white points) to an approximate location in visual space (bottom, white points). RF responses at these locations were estimated as weighted averages of previously simulated RF responses in visual space (bottom, black points).

Simulated RF responses for stimuli with different apertures resulted in qualitatively similar aperture-dependent orientation preferences as seen in the mouse sSC imaging data. For example, consider RF orientation preferences for the smallest circular aperture (Fig. 3a, 20° radius column). In our model and the mouse sSC data, RFs responsive to locations along the stimulus aperture (Fig. 3a, radius 20°, top panel, locations 1-6) demonstrated robust orientation preferences (Fig. 3a, cf. middle & bottom panels, locations 1-6). In contrast, RFs responsive to locations away from the aperture, such as towards the center of the stimulus (Fig. 3a, radius 20°, top panel, location “central”), tended not to exhibit orientation preferences (Fig. 3a, cf. middle & bottom panels, location “central”). Notably, the orientation preferences from both the imaging data and model predictions varied systematically along the aperture: the preferred orientation at each RF location was orthogonal to the aperture edge. We refer to this phenomenon as a preference for *radial orientations*. While this term is sometimes used to denote orientations radial with respect to the fovea^16–18^, here we use it to describe orientations radial to the center of the stimulus. For example, a RF responsive to the *horizontal* (top) edge of the aperture (Fig. 3a, radius 20°, top panel, location 4) preferred a *vertical* orientation (Fig. 3a, cf. middle & bottom panels, location 4, see also Fig. 3a color bar, circle 4, cyan), while an RF responsive to the *vertical* (left) edge of the aperture (Fig. 3a, radius 20°, top panel, location 1) preferred a *horizontal* orientation (Fig. 3a, cf. middle & bottom panels, location 1, see also Fig. 3a color bar, circle 1, red). Likewise, RFs responding to other locations along the aperture (Fig. 3a, radius 20°, top panel, locations 2, 3, 5, 6) also demonstrated orientation preferences orthogonal to their respective aperture edges (Fig. 3a, cf. middle & bottom panels, locations 2, 3, 5, 6; orientation preferences: 15°, 60°, 120°, 150°, see color bar circles 2, 3, 5, 6).

**Figure 3.**
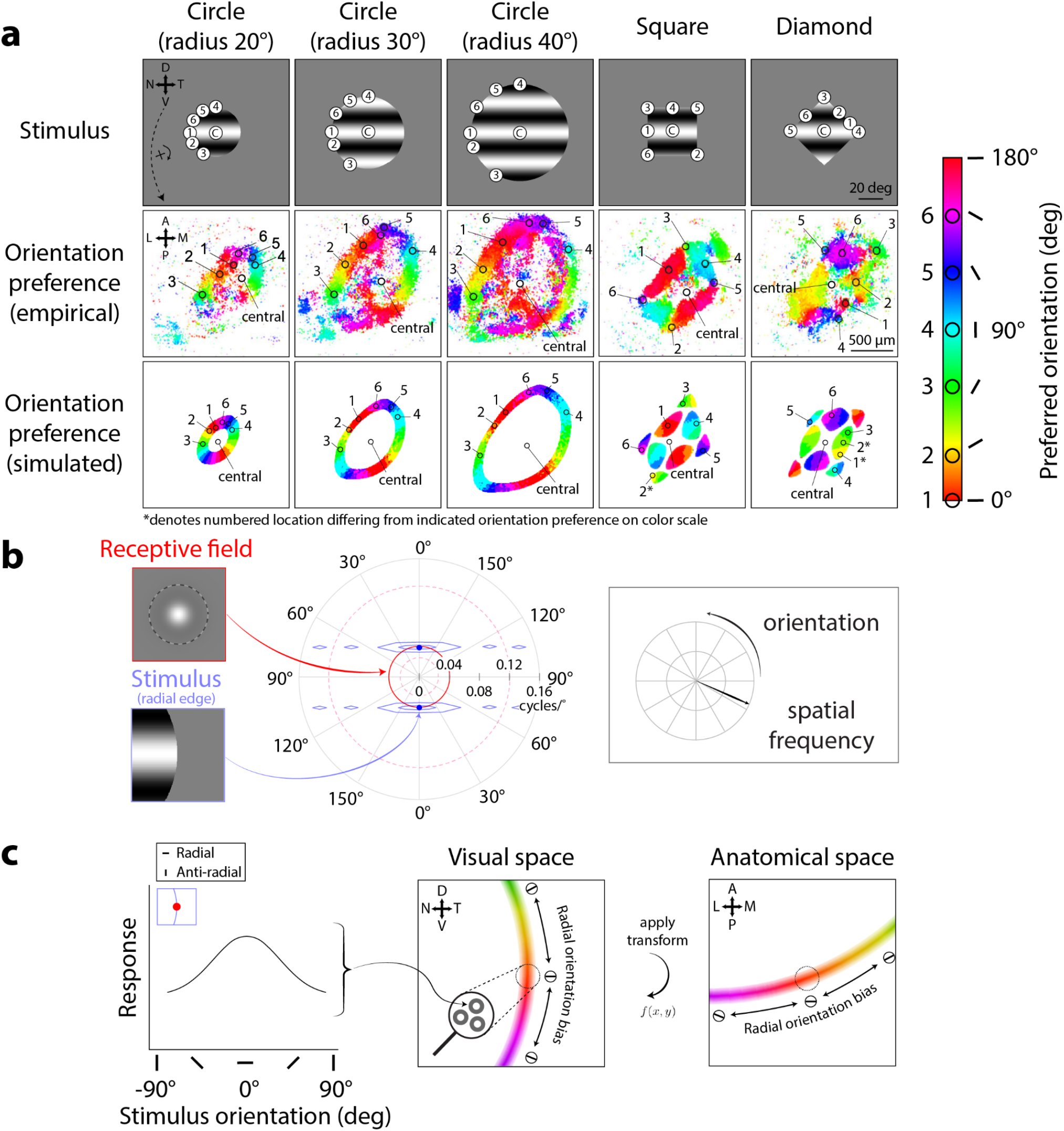
Center-surround RF model predicts orientation preference map changing across aperture size and shape. **A)** Orientation preference maps from Ca^2+^ imaging (middle row, adapted from Fig. 2, Ref. 9) and RF model simulations (bottom row) to oriented grating stimuli of varying aperture shapes and sizes (top row). Map colors correspond to preferred orientation, see scale. Numbered circles in color scale and in the middle row, leftmost panel were defined from prior imaging data^9^ to show varied orientation preferences around SC; we matched these numbered circles onto all other panels to illustrate matching retinotopic and anatomical locations. “C” (top row): central. **B)** Idealized interaction in spatial frequency domain between center-surround RF (red inset) and an edge that is perfectly vertical (blue inset). Overlap between RF (red solid/dashed circles, center) and stimulus (blue envelopes, center) determines RF response magnitude. Solid red circle denotes RF center (preferred) spatial frequency; dashed red circles denote ±1 standard deviation from center spatial frequency in log-units. **C)** Left panel: Model single-unit center-surround RF responses across stimulus orientations at vertical aperture edge (inset). Center and right panels: Population RF orientation preferences (oriented markers and color gradient, same scale bar as in A) from averaging single-unit RF responses across spatial locations (e.g., center panel, zoom-in), shown in visual (center) and corresponding anatomical coordinates (right).

Across aperture sizes, orientation preference maps predicted by the model provided a qualitative match to the maps from imaging data. In response to larger circular apertures (Fig. 3a, radius 30° & 40°, top panels), the circular pattern of RF orientation preferences expanded in both the simulated results and the imaging data. This expansion also resulted in a larger retinotopic region corresponding to the center of the stimulus (Fig. 3a, radius 30° & 40°, top panels, locations around central) lacking orientation preferences (Fig. 3a, radius 30° & 40°, middle & bottom panels, non-colored regions around “central”). Although the locations in the sSC exhibiting orientation preference changed with the size of the aperture, radial orientation preferences remained consistent (Fig. 3a, cf. orientation preference color gradients, i.e., red → orange → yellow → …, across radius 20°, 30°, 40°, middle & bottom panels).

Across aperture shapes, simulated orientation preference maps also qualitatively matched the maps from imaging data. For the square and diamond apertures (Fig. 3a, square & diamond columns, top panels), the previously demonstrated circular patterns of RF orientation preferences now adopted rectangular patterns, shown in both the model and imaging data (Fig. 3a, square & diamond columns, cf. middle & bottom panels). RFs lacked orientation preferences at locations near the aperture center (Fig. 3a, square & diamond columns, top row, locations around central). Along the straight edges of the square, the imaging data demonstrated clear radial orientation preferences (Fig. 3a, square column, middle panel, location 1, orientation preference: 0°, red; location 4, orientation preference: 90°, cyan). This pattern was also evident in our simulation (Fig. 3a, square column, bottom panel, location 1, orientation preference: 0°, red; location 4, orientation preference: 90°, cyan). At corner locations, our model again predicted orientation preferences similar to those measured in the imaging data (Fig. 3a, compare middle and bottom panels, square column, locations 2, 3, 5, 6; diamond column, locations 3, 4, 5). However, we note some difference in the orientation preference maps in response to the diamond aperture; the imaging data showed a more scattered set of orientation preferences along the aperture edge than the model (Fig. 3a, diamond column, compare orientation preferences of locations 1 & 2 between middle & bottom panels).

The reason the center-surround RF model produced radial orientation preferences can be readily appreciated by considering the overlap of the stimulus energy with RFs. We first provide a heuristic explanation in the spatial domain. At any edge of a grating stimulus, the stimulus can be thought of as having two different orientations - the orientation of the grating and the orientation of the edge. For example, at the rightmost edge of the stimulus, a sinusoidal grating vignetted by an aperture contains both the horizontal orientation of the grating, as well as the vertical orientation of the aperture edge (Fig. 3b, blue inset). For a circular-symmetric RF not intrinsically selective for orientation (Fig. 3b, red inset), radial orientations, which primarily have stimulus energy at these two orthogonal orientations, produce a larger RF response compared to “anti-radial” orientations (i.e., stimulus orientations running parallel to edge orientation, see Fig. 4b, bottom row insets for examples) where there is only a single orientation in the stimulus (see^19^ for an in-depth discussion of this phenomenon).

**Figure 4.**
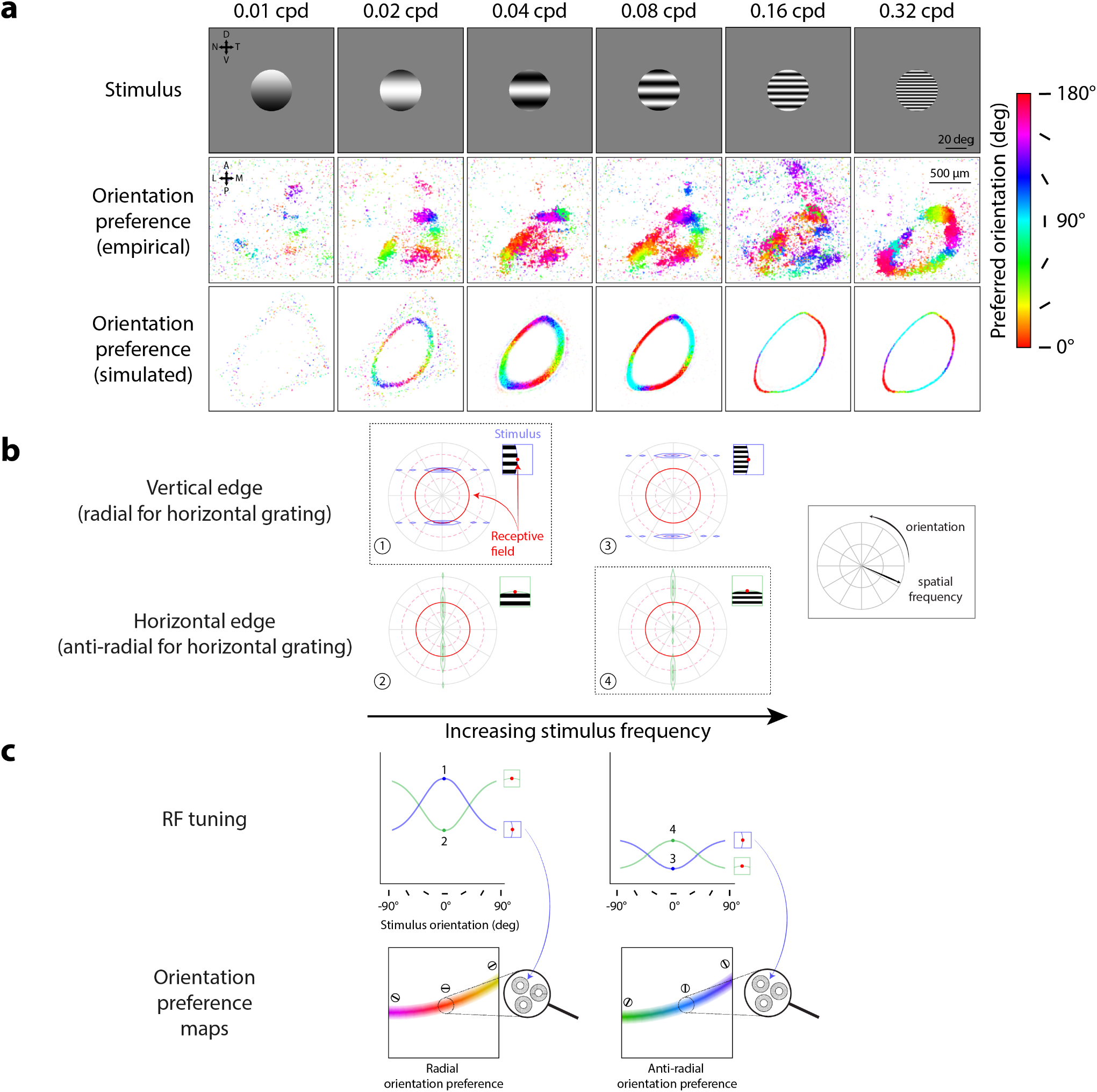
Center-surround RF model predicts orthogonal orientation preferences changing with stimulus spatial frequency. **A)** Orientation preference maps from imaging data (middle row, adapted from Supplementary Fig. 3, Ref. 9) and model simulations (bottom row) in response to oriented grating stimuli of varying spatial frequencies; map colors correspond to preferred orientation, see scale; cpd: cycles/°. **B)** Model prediction of largest RF responses (dotted outlines) depending on overlap between RF (red) and horizontal stimulus grating at vertical (blue, panels 1, 3) or horizontal (green, panels 2, 4) edges. Left to right: increasing stimulus frequency. Solid red circles denote RF center (preferred) spatial frequency; dashed red circles denote ±1 standard deviation from center spatial frequency. **C)** Model prediction of orientation preferences across stimulus spatial frequencies; left to right: low to high spatial frequency. Top row: Orientation tuning curves for single-unit RFs at vertical (blue, see inset) and horizontal aperture edges (green, see inset); numbered points correspond to panels in B (RF responses to horizontal gratings). Bottom row: Orientation preference maps for population RFs across simulated anatomical locations. Dotted circles show example population RFs around the vertical aperture edge, generated by averaging local single-unit RF responses with tuning profiles shown in the top row (blue arrows); color gradient (see scale in A) and oriented markers indicate varying population orientation preferences along the stimulus aperture.

The explanation for radial orientation preferences is more apparent by considering the RF and stimulus in the spatial frequency domain. Because the model center-surround RFs were constructed as a difference-of-Gaussians in the spatial domain, the RFs in the spatial frequency domain were also difference-of-Gaussians^20^. Thus, these RFs formed a donut-like shape in the spatial frequency domain (Fig. 3b, red solid/dashed circles) selective for spatial frequency. To determine how such a linear RF would respond to a stimulus, we can examine the overlap of the stimulus with the RF in the spatial frequency domain. The response is the pointwise product of the RF and the stimulus, summed across all spatial frequencies. For example, consider such an RF with a center spatial frequency of 0.04 cycles/° (Fig. 3b, red inset) that overlaps with the rightmost vertical, i.e., *radial*, edge of a horizontally oriented sinusoidal grating, also with a spatial frequency of 0.04 cycles/° (Fig. 3b, blue inset). An infinitely large grating in the spatial domain is represented by two infinitesimally small points (Fig. 3b, blue points) in the spatial frequency domain, with their distance from the origin determined by the grating’s spatial frequency (e.g., 0.04 cycles/°). For a horizontal grating (0°/180° orientation), the angular locations correspond to 90° and 270° in polar angle. The presence of an aperture edge spreads the stimulus energy across orientations and spatial frequencies, depending on the orientation of the edge. For example, a perfectly vertical aperture edge spreads stimulus energy horizontally, primarily across many orientations, in the spatial frequency domain (Fig. 3b, center, blue envelope). To generalize, any oriented edge will spread stimulus energy in the orthogonal direction in the spatial frequency domain (e.g., a horizontal aperture edge would spread energy vertically). Critically, this means that the *radial aperture edge* will spread stimulus energy primarily across *many orientations*. In the case that the stimulus spatial frequency is near the RF’s center spatial frequency, the stimulus orientation creating a radial aperture edge (in this example, a vertical aperture edge for a horizontal grating stimulus) creates the largest overlap between stimulus and RF (Fig. 3b, blue and red overlap), compared to any other orientation. Therefore, grating orientations that create a radial aperture edge are predicted to generate the largest RF response out of all possible orientations (Fig. 3c, left panel).

Extending this logic to a spatially distributed population of center-surround RFs, we arrive at the model predictions described earlier (Fig. 3a, bottom row). Orientation preference maps changed dynamically based on stimulus shape and size in the imaging data (Fig. 3a, middle row). Our model shows that preferences for radial orientations systematically explain this phenomenon because a spatial distribution of single-unit center-surround RFs preferring radial orientations should produce a gradient of radial orientation preferences varying smoothly along the aperture edge. This can be appreciated intuitively in visual coordinates (Fig. 3c, center) and equivalently in anatomical coordinates (Fig. 3c, right) for comparison to the imaging data. Additionally, anatomical locations representing the center of the stimulus tended to lack orientation preferences (Fig. 3a, middle row, non-colored locations around “central”). This too is predicted by the model, because where there is no spreading of orientation energy (i.e., where RFs do not overlap the stimulus edge), there should be no preferred orientation. Across stimulus shapes and sizes, the logic of stimulus and RF overlap implemented by our model captures the stimulus-dependent nature of orientation preferences found in the imaging data. In summary, linear circular-symmetric RFs are sufficient to account for the stimulus-dependent dynamic orientation phenomena observed in prior imaging data.

### Orientation preference maps depend on stimulus spatial frequency

Our model predicts that radial orientation preferences should change with the spatial frequency of the stimulus, an effect that was also found in mouse sSC data. Orientation preferences in mouse sSC were previously obtained^9^ using a set of stimuli varying in spatial frequency (Fig. 4a, top row), spanning spatial frequency preferences reported in the mouse sSC literature^11,15^. The imaging results showed that the most robust representation of radial orientation preferences appeared for stimuli with spatial frequencies of 0.04 and 0.08 cycles/° (Fig. 4a, 0.04 and 0.08 cycles/° column, middle row). To simulate RFs that could reproduce this phenomenon, we performed a grid search across a range of center-surround RF parameters (center-surround size and amplitude ratios, which vary RF spatial frequency tuning) to find RF parameters producing orientation preference maps that best matched the imaging results qualitatively (see following section and accompanying Fig. 5 for a more detailed description). Center-surround RFs with a size (σ) ratio of 0.8° to 4.7° and surround-to-center amplitude ratio of 0.4 to 1 (resulting in a preferred spatial frequency of 0.08 cycles/°) predicted orientation preferences (Fig. 4a, 0.04, 0.08 cycles/° columns, bottom row) best matching the empirical observations (Fig. 4a, 0.04, 0.08 cycles/° columns, middle row), demonstrating radial orientation preferences for stimuli with spatial frequencies of 0.04 and 0.08 cycles/°. At lower stimulus spatial frequencies, the imaging data demonstrated a fading of the radial orientation preference map (Fig. 4a, 0.01, 0.02 cycles/° columns, middle row), which is also predicted by the model (Fig. 4a, 0.01, 0.02 cycles/° columns, bottom row).

**Figure 5.**
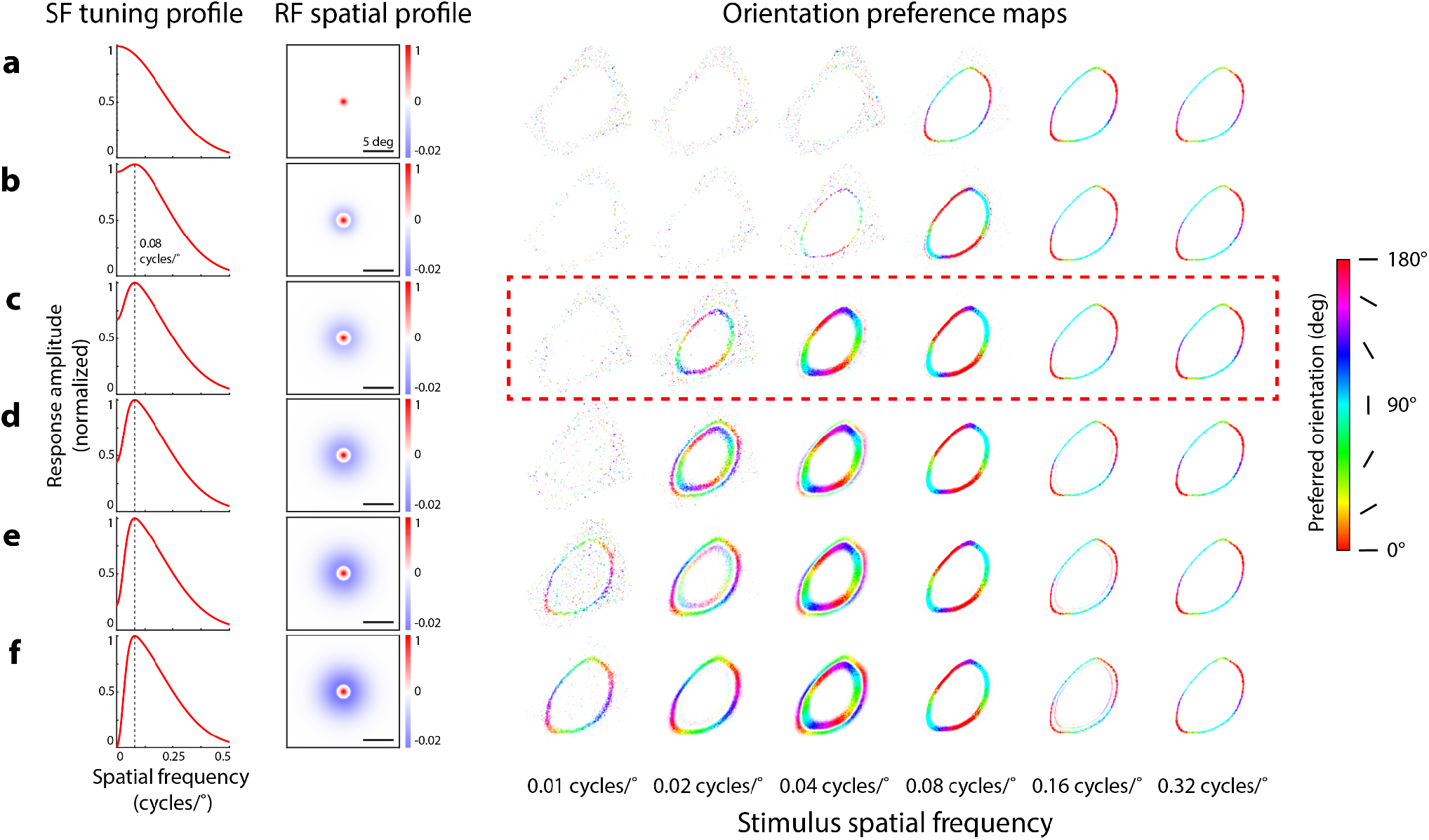
Impact of surround size and amplitude on simulated orientation preference maps. **A)** Simulated orientation preference maps (right) produced by spatially distributed, circular-symmetric Gaussian RFs (center size, σ: 0.8°) in response to circular grating stimuli with a radius of 30°. Colors correspond to orientation preference (color scale, right). **B-F)** Simulated orientation preference maps as described in A, using center-surround RFs with varying center-surround size (σ) ratios and center-surround amplitude ratios to maintain a preferred spatial frequency (SF) of 0.08 cycles/° (dashed vertical lines, left column) across all RF manipulations. Dashed red box corresponds to the same orientation preference maps shown in Fig. 4. Center to surround size ratios (B-F): 0.8° to 3, 4.7, 5.2, 5.5, 5.7°; center to surround amplitude ratios: 1 to 0.2, 0.4, 0.6, 0.8, 1. Responses in SF tuning profiles normalized from 0 to 1, after thresholding and summing responses from pairs of opposite polarity center-surround RFs (see Methods). Amplitudes of RF spatial profiles were normalized between possible values of −1 to 1 (scale shown from −0.02 to 1 for clearer visualization of On-center RFs, paired Off-center RFs not shown). Amplitude ratio refers to the ratio of volumes under the circular 2D center and surround Gaussians used to construct the difference-of-Gaussians center-surround RFs.

Notably, prior imaging data revealed that at high stimulus spatial frequencies, orientation preferences shifted from radial orientation preferences to *anti-radial* orientation preferences, a result that our center-surround RF model also captured. We refer to anti-radial orientations as orientations parallel to the edge, i.e., orthogonal to radial orientations. Anti-radial orientation preferences were evident for the highest stimulus spatial frequency in both the imaging data (Fig. 4a, middle row, 0.32 cycles/°) and the model-predicted orientation preference maps (Fig. 4a, bottom row, 0.32 cycles/°).

The emergence of anti-radial rather than radial orientation preferences can be explained by our model when considering the spatial frequency representations of stimuli with frequencies higher than the RF’s preferred spatial frequency (Fig. 4b, right column). For example, consider a horizontal stimulus grating (Fig. 4b, insets). As stimulus spatial frequency is increased, stimulus energy in the spatial frequency domain is shifted outward, meaning that stimuli with higher spatial frequency are now pushed towards the outer edge of the RF (Fig. 4b, compare stimulus energy locations between left and right columns). A vertical (radial) edge spreads stimulus energy across a much smaller portion of the RF at high spatial frequencies (Fig. 4b, panel 3), compared to when stimulus and RF preferred spatial frequencies are matched (Fig. 4b, panel 1). In contrast, the effect of increasing spatial frequency is less consequential when considering stimulus-RF overlap for a horizontal (anti-radial) edge because stimulus energy is spread towards the center of the spatial frequency domain (Fig. 4b, bottom row, compare stimulus-RF overlap between panels 2 and 4). Because the anti-radial aperture edge now spreads a larger amount of stimulus energy across the RF than the radial aperture edge (Fig. 4b, right column, compare stimulus-RF overlap between panels 3 and 4), we observe a larger RF response at the anti-radial over the radial aperture edge (Fig. 4c, right column, top panel, compare responses at points 3 and 4). Averaging across responses from local single-unit RFs, the model predicts a gradient of anti-radial orientation preferences across the population along the aperture edge (Fig. 4c, right column, bottom panel).

A divergence between our model and imaging data in the transition from radial to anti-radial orientation preferences highlights how our model RFs might differ from populations in mouse SC. For the 0.16 cycles/° stimulus, the imaging data appeared to be undergoing a somewhat messy, non-uniform transition between radial and anti-radial orientation preferences (Fig. 4a, 0.16 cycles/° column, middle panels), but our model produced a clear map of anti-radial orientation preferences. The difference between the two could be due to several factors. Since the model predicted a transition between 0.08 and 0.16 cycles/°, the modeled population RFs’ spatial frequency tuning profile may not have exactly matched those in mouse sSC. Additionally, the model’s simplified assumption of uniform RF properties across mouse sSC may have played a role as well. Population RFs pooled from many single units with a range of different RF properties may better recapitulate the less uniform transition from radial to anti-radial orientation preferences across stimulus spatial frequencies.

### Changes in orientation preference maps depend on RF parameters

Simulations of different center-surround RF properties revealed that small changes in RF properties can alter the transition between radial and anti-radial orientation preference maps. In a grid search procedure, we generated orientation preference maps to stimulus spatial frequencies ranging from 0.01-0.32 cycles/° using RFs varying in center-surround size and amplitude ratios while constrained to a particular preferred spatial frequency (Fig. 5a-f, SF tuning profile, RF spatial profile columns). As surround amplitude increased, we found the emergence of anti-radial orientation preferences at low spatial frequencies (Fig. 5e-f, orientation preference maps, 0.01 and 0.02 cycles/° columns). Higher surround amplitudes also resulted in orientation maps with inner and outer rings of opposing orientation preferences (Fig. 5d-f, orientation preference maps, 0.02 and 0.04 cycles/° columns). Notably, model RFs without a surround were unable to produce maps showing radial orientation preferences (Fig. 5a, orientation preference maps).

While small differences in RF properties changed where the radial to anti-radial transition point occurred, all modeled center-surround RFs showed a transition to anti-radial orientation preferences at higher stimulus spatial frequencies, similar to the imaging results from mouse SC.

We found that maps of anti-radial orientation preferences produced in response to high spatial frequency stimuli were robust across our manipulations of RF parameters (Fig. 5a-f, orientation preference maps, 0.16 and 0.32 cycles/° columns). Similar patterns of results were found for RFs constrained to other preferred spatial frequencies (0.02, 0.06, 0.1 cycles/°, not shown). The only notable difference caused by a change in preferred spatial frequency was the shifting of stimulus spatial frequency where the transition from radial to anti-radial orientation preferences was observed. Altogether, we found that orientation preference maps were sensitive to manipulations of surround amplitude and size, even while maintaining the same preferred spatial frequency. Our results suggest that the overall shape of the spatial frequency tuning curve, not just the peak tuning, dictates how patterns of orientation preferences manifest in sSC populations.

### RF structures from single unit and population-scale measurements

The center-surround RF model not only behaved similarly to population data in mouse SC, but also captured orientation preferences for single units measured with narrow-field imaging. Center-surround RFs in our model simulations produced single-unit orientation preferences (Fig. 6a, center column, middle panel) that qualitatively matched the single-unit imaging data (Fig. 6a, left column, middle panel), demonstrating radial orientation preferences dependent on edge orientation (Fig. 6b, c, left column, top panel). These simulated single-unit preferences were maintained at the population scale (Fig. 6b, left column, bottom panel), since the population response at each spatial location was simply obtained by averaging responses across nearby units (Fig. 6c, left column, bottom panel, dotted circle).

**Figure 6.**
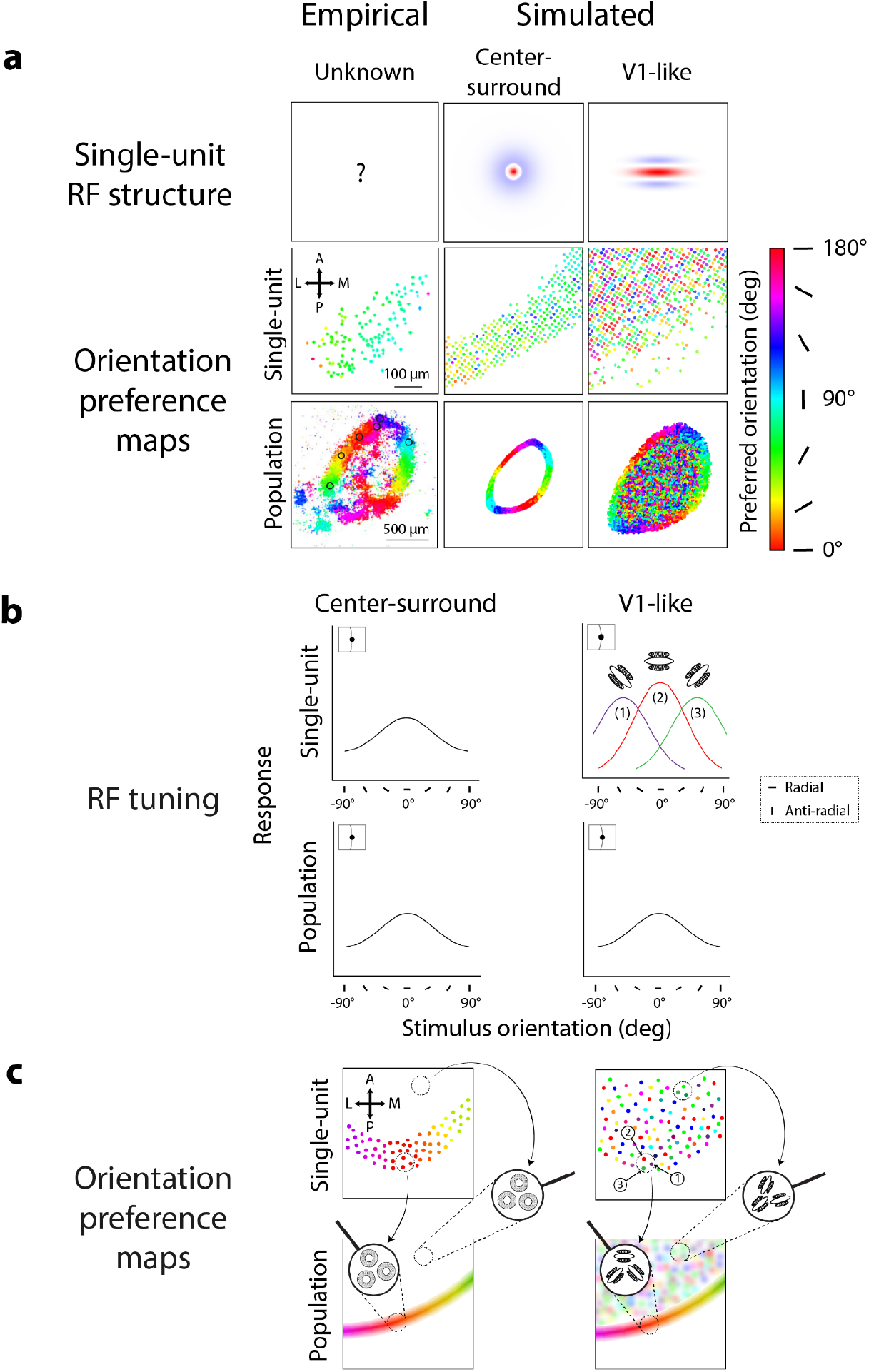
Center-surround and V1-like models produce different patterns of orientation preferences at population and single-unit scales. **A)** Orientation preference maps from center-surround and V1-like RFs (top row) at single-unit (middle row) and population scales (bottom row) in response to circular grating stimuli (spatial frequency: 0.04 cycles/°, radius: 30°); colors correspond to orientation preference, see scale. Empirical maps (middle & bottom panels, left column) adapted from Figs. 2 & 3, Ref. 9. **B)** Model-predicted orientation tuning for center-surround and V1-like RFs at single-unit and population scale, shown for RFs located at the vertical aperture edge (insets). Color in top right panel corresponds to preferred orientation for example V1-like RFs 1-3 (see scale in A). Legend denotes radial and anti-radial orientations for vertical aperture edges. **C)** Model-predicted orientation preference maps for center-surround and V1-like RFs at single-unit and population scale for RFs distributed across SC. Responses from nearby single-units (top panels, dotted circles) are combined to generate local population orientation preferences (bottom panels, zoom-ins). Numbered units in top right panel correspond to example single V1-like units shown in B; colors correspond to preferred orientation (see scale in A).

Intrinsically orientation-selective V1-like RFs showed different orientation preferences at the single-unit scale compared to the imaging data and the center-surround RF simulations. Here, we define a V1-like RF as an RF with elongated subfields in parallel, as originally identified in V1 of cat and macaque^21,22^. We simulated V1-like RFs as oriented Gabor filters^10,23–26^ (Fig. 6a, top row, right column), with the same preferred spatial frequency as the simulated center-surround RFs (0.08 cycles/°) and with a random orientation at each spatial location. Simulated individual V1-like RFs thus had orientation preferences that varied randomly across model sSC (Fig. 6a, right column, middle panel), in contrast to the smooth gradient of orientation preferences in both the single-unit imaging data (Fig. 6a, left column, middle panel) and center-surround simulation (Fig. 6a, middle column, middle panel). Orientation preferences for simulated V1-like single units were scattered randomly because responses to different stimulus orientations depended largely on the intrinsically oriented RF structure (Fig. 6b, right column, top panel), making it so the stimulus aperture had minimal influence on orientation preferences (Fig. 6c, right column, top panel, example RFs 1-3 along stimulus edge).

V1-like RFs could, however, produce radial orientation preferences at the population scale. Simulated V1-like RFs at the population scale showed radial orientation preferences along the stimulus aperture (Fig. 6a, right column, bottom panel), as in the imaging data (Fig. 6a, left column, bottom panel) and the center-surround RF simulations (Fig. 6a, middle column, bottom panel). We note a larger overall region of orientation-selective responses from V1-like RFs compared to center-surround RFs (Fig. 6a, bottom row, middle vs. right panels) due to the larger size of the individual V1-like RFs. Radial orientation preferences for V1-like populations emerged because of the pooling of responses from randomly oriented V1-like single units. In theory, if each local population averages responses from units of all orientation preferences, the population RF will no longer be tuned for orientation and thus only be selective for spatial frequency, similar to center-surround RFs. This center-surround-like population RF is thus susceptible to aperture-dependent radial orientation preferences^27,28^ (Fig. 6b, right column, bottom panel). Therefore, along the stimulus aperture, population RFs composed of randomly oriented V1-like RFs should demonstrate radial orientation preferences (Fig. 6c, right column, bottom panel, left zoom-in). Without the influence of the stimulus aperture, these population RFs should have no orientation preference, like the center-surround RFs, yet our simulations demonstrated a patchy pattern of orientation preferences within the stimulus aperture (Fig. 6a, right column, bottom panel). Within the aperture, orientation preferences arose from local regions of V1-like single units over-representing certain intrinsic orientation preferences over others^29,30^ (Fig. 6c, right column, top panel, right zoom-in). Thus, the resulting population RF maintained some intrinsic orientation preference determined by the preferences of its underlying single units (Fig. 6c, right column, bottom panel, right zoom-in).

### Preference-maintaining mouse sSC neurons are better modeled as V1-like RFs

Imaged mouse sSC neurons showed a mixture of orientation preference-switching and preference-maintaining properties, which could be modeled by a proportion of V1-like RFs intermingled with center-surround RFs. In the Ca^2+^ imaging data, responses were obtained from a set of 53 neurons at single-cell resolution to oriented gratings vignetted by a horizontal (Fig. 7a, left) and vertical aperture (Fig. 7b, left), as well as with no aperture (Fig. 7c, left). These neurons had RFs centers located within the central 20 degrees of the visual field (Fig. 7a-c, left column, green boxes). The empirical results showed that the set of neurons largely preferred vertical orientations when presented with a horizontal aperture (Fig. 7a, middle-left), horizontal orientations when presented with a vertical aperture (Fig. 7b, middle-left), and random orientations when no aperture was present (Fig. 7c, middle-left). We presented the same stimuli to simulated RFs with RF centers within the same region of visual space and in the same relative spatial configuration as in the imaging data (Fig. 7a-c, middle-right & right columns). Simulated center-surround RFs reproduced the preference-switching effect shown in the imaging data (Fig. 7a, b, middle-right column) and produced random idiosyncratic orientation preferences due to simulated noise in the absence of stimulus apertures (Fig. 7c, middle-right column). In contrast, simulated V1-like RFs maintained their orientation preferences across aperture conditions (Fig. 7a, b, right column) and did not have their orientation preferences changed by the same amount of simulated noise (Fig. 7c, right column).

**Figure 7.**
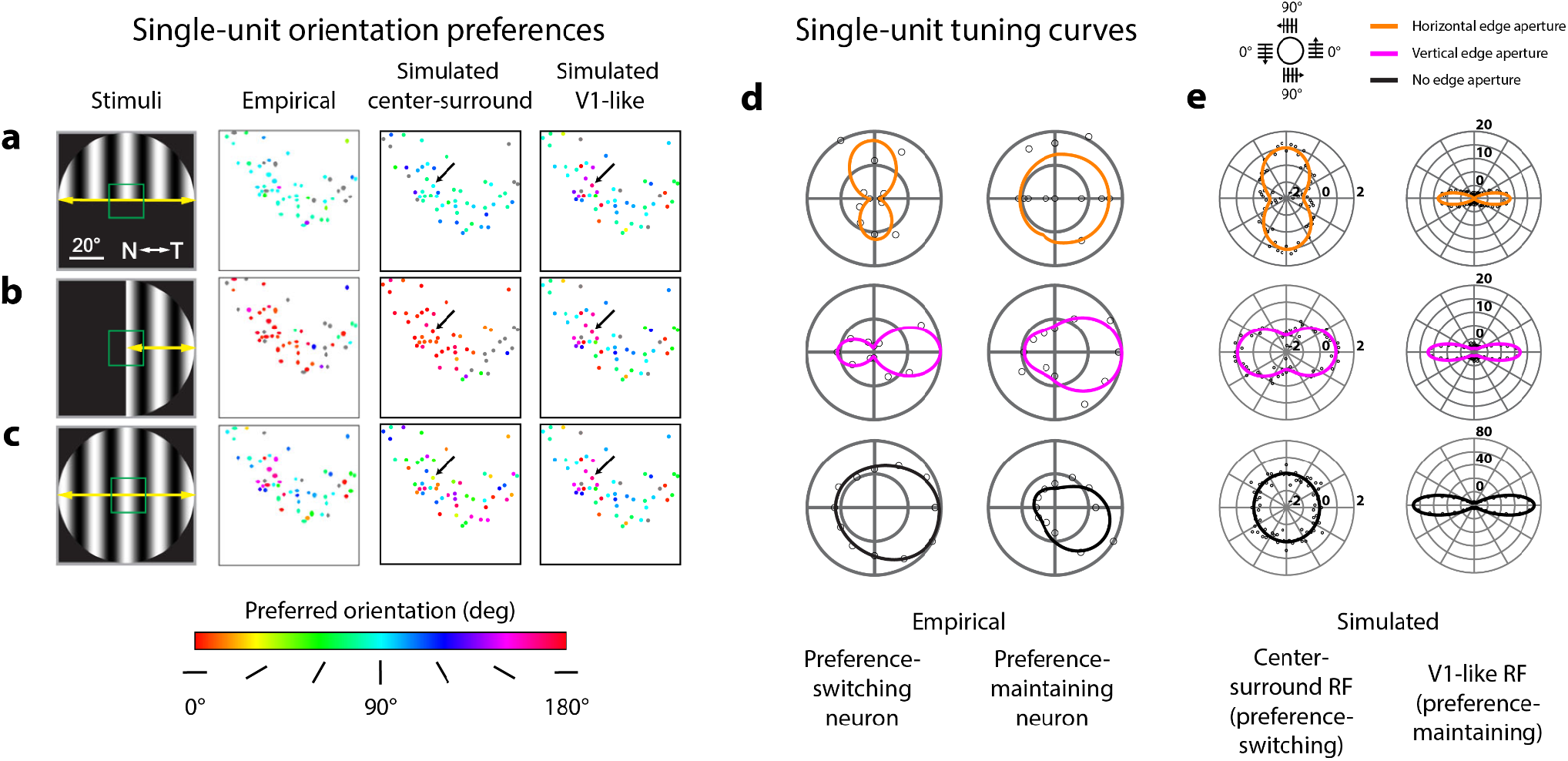
Center-surround and V1-like RF structures produce preference-switching and preference-maintaining tuning patterns. **A)** Maps of orientation preferences measured empirically from single neurons (middle-left column), simulated using center-surround RF structure (middle-right column), and simulated using V1-like RF structure (rightmost column), in response to grating orientations vignetted by a horizontal edge (leftmost column). Colors correspond to preferred orientation (color scale below panel C). RFs from empirical and simulated neurons fall within the green box shown in visual space (leftmost column); corresponding anatomical locations in either physiological or simulated sSC are depicted in the single-unit maps. Left and middle-left columns adapted from Fig. 4A-C, Ref. 9. **B)** Orientation preferences as described in A, but with grating stimuli vignetted by a vertical edge. **C)** Orientation preferences as described in A, but without any edge aperture. **D)** Empirically measured responses from example preference-switching (left column) and preference-maintaining (right column) neurons to drifting grating orientations (polar plot legend above panel E) vignetted by horizontal, vertical, or no edge apertures (color legend above panel E). Data and visualization adapted from Fig. 4E-F, Ref. 9. **E)** Simulated responses from example center-surround (left column) and V1-like (right column) RFs to static grating orientations, vignetted by horizontal, vertical, or no edge apertures. Simulated center-surround responses (left column) were generated by the highlighted center-surround unit (A-C, middle-right column, black arrows); simulated V1-like responses (right column) were generated by the highlighted V1-like unit (A-C, rightmost column, black arrows). Simulated responses from 0-180° grating orientations were duplicated for 180-360° to match Fig. 7D. Units on polar plots correspond to percent change from the mean response across all orientations from the indicated neuron (black arrows, A-C).

Additionally, we matched the orientation tuning shapes of two types of neurons described in single-unit Ca^2+^ imaging using simulated center-surround and V1-like RFs. Prior imaging data had revealed examples of neurons that either switched their orientation preference (Fig. 7d, left column) or largely maintained their orientation preference (Fig. 7d, right column) in response to stimuli of different aperture types. We found that simulated center-surround single-unit RFs produced similar shifts in tuning as the empirically measured preference-switching neurons, in response to apertures of different orientations (compare Fig. 7d and 7e, left columns). For simulated V1-like RFs, we found that orientation preferences stayed consistent for the same orientation across aperture types, with their orientation tuning being dictated mainly by the inherently oriented structure of the RF (Fig. 7e, right column). These V1-like RFs matched the behavior of the empirically measured preference-maintaining neurons (compare Fig. 7d and 7e, right columns).

There were also several discrepancies between the tuning shapes from the imaging and simulated data. We note that the empirically-derived tuning curves showed a substantial degree of direction selectivity (Fig. 7d, note tuning asymmetry). Our simulations aimed to explain how purely spatial changes in RF properties can result in changing orientation preferences. Thus we explicitly modeled spatial RFs without a temporal component, which cannot capture response differences from motion direction (see Discussion). Another discrepancy was the width of the tuning, or degree of orientation selectivity, between the empirical and the simulated tuning curves, particularly for V1-like RFs (Fig. 7d, e, right columns). The degree of orientation selectivity was affected by the aspect ratio of the simulated V1-like subregions, and our simulated subregion aspect ratios were based on measurements previously reported in cat striate cortex^10^. Further empirical evidence may provide better estimates for what values should be expected for mouse SC.

In summary, these results suggest that center-surround RFs are likely prevalent in mouse sSC as demonstrated by the substantial proportion of neurons shifting their orientation preference in the imaging data. However, the presence of single units that maintained their orientation preference regardless of the aperture suggests a proportion of neurons with V1-like RF structure are intermingled among a larger population with center-surround RF structure.

## Discussion

We show that a simulation based on classic RF models can recapitulate stimulus-dependent changes in population orientation selectivity in mouse sSC^9^. We found that simulated orientation preference maps changed with stimulus shape and size, reproducing a wide range of Ca^2+^ imaging results without requiring changes in selectivity. A prediction of our model is that population orientation selectivity should change from radial to anti-radial with increases in spatial frequency, which is consistent with the Ca^2+^ imaging data. We found that both center-surround and V1-like RFs each captured key elements of the empirical data^9^. Narrow-field Ca^2+^ imaging at single-cell resolution identified some neurons that maintained orientation preference, while others switched preference with changes in the stimulus aperture. Our modeling results suggest this may be attributable to differences in RF structure between these two subpopulations. The preference-switching subpopulation was better captured with center-surround structure and the preference-maintaining subpopulation with V1-like structure. As primate SC neurons are not traditionally thought to have orientation-selective, V1-like RFs^31–33^ (but see^34,35^), this suggests that the preference-maintaining subpopulation represents a fundamental difference between rodent and primate visual systems. Nonetheless, our modeling demonstrates that while RF properties of some neurons in mouse sSC may differ from those in primate SC, all neurons conform to canonical computations that have been well-established in early parts of the visual pathway^20,36– 38^.

Our results demonstrate that rewiring of RFs is not required to account for stimulus-dependent changes in orientation selectivity. Orientation selectivity in V1 arises from the precise alignment of feedforward projections of center-surround RFs from LGN to V1, creating elongated RFs^1,3,4^. This arrangement is thought to be stable following neural development, robust even to dramatic changes such as retinal lesions^39^. While recurrent feedback within a circuit^40^ may change properties like response gain^41^ or sharpen tuning^42^, feedback is not thought to change the intrinsic stimulus preference of a neuron. Given these classic findings, stimulus-dependent changes in population orientation selectivity in mouse sSC might suggest that mouse anatomical connectivity can change instantaneously. But, by demonstrating that changing population orientation selectivity is predicted by fixed RFs, our results reconcile the mouse sSC findings with classic neurophysiological results, that orientation selectivity arises from stable, feedforward circuitry. This conclusion aligns with theoretical studies^43,44^ explaining apparent shifts in spatial selectivity with attention^45,46^ and eye movements^47–49^, without requiring changes in anatomical connectivity.

While our results demonstrate that center-surround RFs are sufficient to produce population orientation selectivity, two lines of evidence suggest that a subpopulation of neurons in mouse sSC is inherently orientation selective. First, a number of studies have reported orientation selectivity in mouse retinal ganglion cells (RGCs)^50–52^, a property that may arise from dendritic morphology^52^ of genetically identified RGC subtypes (e.g., JAM-B RGCs^53^). The proportion of RGCs that project to sSC is far higher in mice (85-90%)^54^ than cats (~50%)^55^ or primates (~10%)^56^, so inputs to mouse sSC likely include those from orientation-selective RGCs. Removal of V1 does not generally affect orientation-selective patterns in mouse sSC^9,11,57,58^, suggesting that mouse V1 is likely not the source of orientation selectivity in mouse sSC. Second, studies using white noise stimuli and reverse correlation analyses, methods that are considered to be unbiased by stimulus edges^59^, have revealed oriented RF structure in mouse retina (ON and OFF OS RGCs)^50,60^, LGN (relay cells)^50,61^, and mouse sSC (Ntsr1+ cells)^62^. Our modeling approach provides a means of distinguishing the degree to which population orientation selectivity arises from intrinsically oriented RFs, center-surround RFs, or a combination of both.

An extension of the logic of our model to the temporal domain might allow for a similar description of population direction selectivity. The model implemented here used purely spatial RFs designed to explain spatial response properties and therefore cannot explain temporal properties, such as direction selectivity in mouse sSC^9,11,12,14,58,62–64^. Not only would adding a temporal dimension to our model allow for the possibility of explaining direction selectivity, but may also provide an explanation for other unintuitive results, such as changes in orientation selectivity resulting from a wider range of stimulus-dependent properties. For example, a prior study demonstrated how a spatiotemporal RF model of V1^65^ can produce apparent orientation tuning shifts in ferret V1 population responses depending on stimulus size, speed, and direction^66^. Adding a temporal component would incorporate a range of findings characterizing temporal tuning properties of mouse sSC neurons^9,11,12,14,58,62–64^ which may provide a more complete explanation of stimulus-dependent selectivity.

The logic we have employed here is analogous to previous studies that have sought to explain orientation selectivity in BOLD fMRI measurements of human V1. Neural responses from individual fMRI voxels reflect the pooled activity of hundreds of thousands of neurons. Even though orientation-selective neurons are organized in cortical columns^1^, forming the well-known pinwheel motif spanning early visual cortex^67^, the scale of the organization is such that a typical fMRI voxel (2.5×2.5×2.5 mm) samples from around 25 orientation columns with a range of orientation preferences^68^. Thus, orientation selectivity is thought to largely cancel out over the population within a single voxel^27,28^, resulting in voxel population RFs that are circular symmetric^69^ and not intrinsically orientation selective. Yet, fMRI responses in human V1 exhibit the same stimulus-dependent orientation selectivity observed in mouse superior colliculus^9^, producing orientation maps that appear radial when the grating stimulus is vignetted by a circular aperture^18^. This stimulus-dependent orientation selectivity has been explained with a computational model constructed with population RFs tuned for spatial frequency, but untuned for orientation^28^, similar to the approach reported here in simulating mouse sSC populations.

A lesson learned from the human fMRI literature is that the dominant component of population responses can be explained by circular symmetric population RFs that are not selective for orientation^28^. If there are any other intrinsically orientation-selective components that contribute to population fMRI responses, a careful modeling approach, an extensive stimulus set of natural images, and a massive dataset are required to parse them out. And even when those components can be measured reliably, they account for only a small proportion of the total variance^70^. These studies suggest that the primary driver of orientation selectivity in fMRI responses in human V1 is the aperture edge, not intrinsic orientation tuning. A very small proportion of the responses may be due to populations with an inherent radial orientation bias^16,71,72^, analogous to the preference-maintaining neural subpopulation in mouse sSC, as the modeling analysis in this current paper suggests.

Parsimonious models grounded in interpretable, mechanistic explanations are indispensable for understanding unifying computational principles shared across model species, though such computations can be implemented in different ways across species. The growing popularity of applying increasingly complex computational models^73,74^ in a range of model systems may capture more variance in the data. But this increase in variance explained can come at the cost of interpretability, i.e., clarity in *how* such models fit the data. The framework we provide here demonstrates the utility of classic models built with interpretable components from fundamental principles of neural computation, as these models provide us with strong, general intuitions for how we expect neural systems to operate.

## Methods

### Overview of simulations

We modeled several simulations with a range of stimuli and with two different RF structures to highlight how phenomena regarding orientation selectivity found in prior imaging results^9^ could be captured by models of circular center-surround and elongated subfield (V1-like) RFs. Simulated neurons with center-surround and V1-like RF structure tiled a grid of coordinates in visual space (Fig. 1a). Simulated responses from these RFs were obtained (Fig. 1c) by presenting simulated RFs with grating stimuli that matched those used to obtain prior imaging data^9^ (Fig. 1b). The grid of RF responses in visual coordinates was then used to interpolate RF responses in anatomical coordinates, matching the coordinate space of the imaging data (Fig. 1d). We simulated population RF responses by applying a spatial smoothing procedure on our simulated single-unit RF responses in anatomical coordinates (Fig. 1e). The main simulations we conducted are briefly described here, with details of simulation and stimuli presented in the sections below. All simulations were performed using code written in Matlab 2023b and are available at https://github.com/austinchkuo/mousesc-ori-model.

#### *Aperture shape simulation* (Fig. 3): Manipulation of stimulus aperture shape and size

Prior imaging experiments^9^ measured orientation-selective responses across mouse sSC neural populations to stimuli with different aperture shapes and sizes. We modeled these responses in simulation using circular-symmetric, center-surround RFs which had no intrinsic orientation preference.

#### *Spatial frequency simulation* (Fig. 4 & 5): Manipulation of stimulus spatial frequency

Prior imaging experiments^9^ measured orientation-selective responses across mouse sSC neural populations to grating stimuli varying in spatial frequency. We modeled these responses in simulation using circular-symmetric, center-surround RFs. We performed a grid search to find center-surround RF parameters providing the best qualitative fit the imaging data (Fig. 5).

#### *RF properties simulation* (Fig. 6 & 7): Manipulation of RF structure at single unit and population scales

Prior imaging experiments^9^ measured orientation-selective responses to grating stimuli with horizontal, vertical, and no edge apertures, both at the population scale and in single sSC neurons. We simulated RFs with center-surround structure and with elongated subfields (V1-like RFs) to compare changes in predicted orientation preferences for both simulated RF types at single-unit and population scales.

### RF models (Fig. 1a)

#### Center-surround RFs

Center-surround RFs were constructed by taking the difference of two, 2D circular-symmetric Gaussian functions in the spatial domain, resulting in a difference-of-Gaussians RF, defined by the following equation:

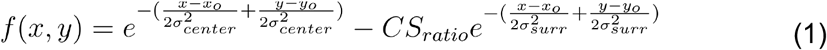

where *x*_*o*_ and *y*_*o*_ denote the center locations of the Gaussians, *σ*_*center*_ and *σ*_*surr*_ denote the standard deviation, or width, of the center and surround Gaussians, and *CS*_*ratio*_ denotes the ratio of surround to center amplitude, specifically the volume under the surround Gaussian to the volume under the center Gaussian. In the aperture shape simulation (Fig. 3), *σ*_*center*_ was 1.5°, *σ*_*surr*_ was 11.4°, and *CS*_*ratio*_ was 0.8. In the spatial frequency simulation and RF properties simulation, *σ*_*center*_ was 0.8°, *σ*_*surr*_ was 4.7°, and *CS*_*ratio*_ was 0.4. We systematically studied the effect of changing these parameters in the grid search corresponding to the spatial frequency simulation (Fig. 5). The values used were *σ*_*center*_ = 0.8, *σ*_*surr*_ = {3, 4.7, 5.2, 5.5, 5.7}, and *CS*_*ratio*_ = {0.2, 0.4, 0.6, 0.8, 1}. Additionally, the grid search also included a condition with no surround, corresponding to *σ*_*center*_ = 0.8, *CS*_*ratio*_ = 0.

A pair of difference-of-Gaussians functions with inverted polarity was created for each location to simulate both On-center and Off-center RFs. The responses from the On-center and Off-center RFs were subsequently thresholded and summed to obtain a single On/Off pair response at each RF location (Fig. 1c).

For the aperture shape simulation, we modeled RFs with a spatial frequency preference of 0.04 cycles/° (*σ*_*surr*_ = 11.4°, *σ*_*center*_ = 1.5°, *CS*_*ratio*_ = 0.8), following RF sizes and spatial frequency preferences reported in the literature^11,15^. In all subsequent simulations, we used center-surround RFs, with *σ*_*center*_ = 0.8°, *σ*_*surr*_ = 4.7°, and *CS*_*ratio*_ = 0.4, following a grid search (Fig. 5) for center-surround RF parameters that produced the best qualitative match to empirical data in the spatial frequency simulation (Fig. 4). These parameters yielded RFs with a center (i.e., preferred) spatial frequency of 0.08 cycles/°. This spatial frequency preference was determined by slicing the Fourier representation of the RF along an axis through the origin to obtain a 1-dimensional Fourier spectrum and recording the spatial frequency corresponding to the peak amplitude.

To simulate the spatial distribution of RF centers across mouse sSC, we created a densely tiled grid of RF locations in visual field coordinates. RF center coordinates *x*_*o*_and *y*_*o*_ranged from −30° to 30°, in 0.25° steps (58,081 RF locations) when calculating responses to circular aperture stimuli with radii of 20°, as well as square and diamond aperture stimuli (e.g., Fig. 3a, top row, far left, center right, far right). We extended the range of both *x*_*o*_ and *y*_*o*_to ±40° (103,041 RF locations) and ±50° (160,801 RF locations) when calculating responses to stimuli that had radii of 30° (e.g., Fig. 3a, top row, center left) and 40° (e.g., Fig. 3a, top row, center).

When simulating preference-switching and preference-maintaining neurons (Fig. 7), we matched 53 simulated single-unit RF locations to the 53 single-unit locations reported in the imaging data, with RF centers falling within the central 20° x 20° square in visual space (Fig. 7a-c, left column, adapted from Fig. 4a-c, Ref. 9). In simulation, the distribution of these RF center locations in anatomical space corresponded to approximately −2.5° to −6.5° from the center of visual space in both X and Y dimensions.

RFs and stimuli were simulated in a 200 × 200 pixel grid representing a −60° to 60° field of view (FOV). Although the pixel sampling may appear sparse, the maximum preferred spatial frequency of our simulated RFs was only 0.08 cycles/°, and the highest stimulus spatial frequency used was 0.32 cycles/°, both of which were well below the Nyquist frequency of 0.83 cycles/° for this FOV.

#### Elongated subfield (V1-like) RFs

V1-like RFs were constructed by pointwise multiplying a 2D sinusoidal grating with an elongated 2D Gaussian envelope. The 2D grating was defined by the following equation,

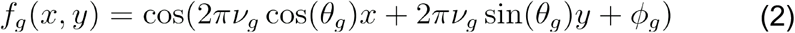

where ν_*g*_ denotes the spatial frequency of the grating in cycles/°, *θ*_*g*_ denotes the orientation of the grating in radians, *ϕ*_*g*_ denotes the phase of the grating in radians.

Note that for the intuitive convention of defining a horizontal grating as 0° and increasing angle as a counterclockwise rotation (thus aligning orientation with polar angle), the following conversion was made,

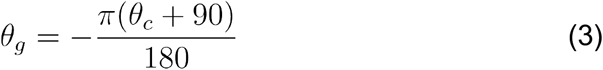

where *θ*_*c*_ denotes orientation (in degrees) in the described convention. Throughout this study, all descriptions of grating orientation are made with respect to *θ*_*c*_. The shape of the Gaussian envelope was defined as follows,

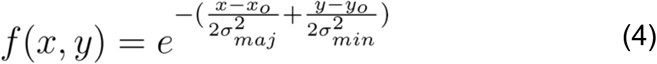

where *σ*_*maj*_ and *σ*_*mim*_ denote the standard deviations of the major and minor axes, and *x*_*o*_and *y*_*o*_denote the location of the RF center. The values of the envelope were then rescaled to range between 0-1. The lengths of the minor and major axes were defined as follows,

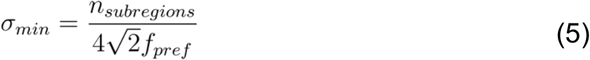

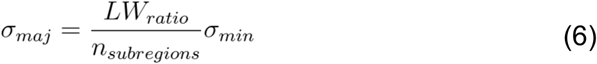

where *n*_*subregions*_ refers to the number of excitatory plus inhibitory subregions in the RF, *f*_*pref*_ refers to the optimal stimulus frequency for the RF (by definition, the frequency of the grating), and *LW*_*ratio*_ refers to the ratio of the major to minor axis lengths. Here, we chose *f*_*pref*_ to be 0.08 cycles/°, to match the preferred spatial frequency obtained from the grid search described for the center-surround RFs (Fig. 5). Prior modeling work of RF properties in cat striate cortex has suggested estimates of *n*_*subregions*_ and *LW*_*ratio*_ as 2 and 4 to be reasonable assumptions^10^. The Gaussian envelope was rotated along with the grating so that the major axis was always aligned parallel to the grating.

The spatial tiling of V1-like RFs was defined by RF center coordinates *x*_*o*_ and *y*_*o*_ ranging ±50°, in 0.25° steps (160,801 RF locations). At each RF location, 8 oriented RFs across equally spaced orientations (*θ*_*c*_ = 0,22.5,45,…,157.5^°^) were constructed. For each RF orientation, we generated two RFs in quadrature phase (*ϕ*_*g*_= 0,90^°^), which were later used to obtain phase-invariant responses. Such responses simulated those in complex cells by squaring and summing responses of quadrature pair RFs^37^ (Fig. 1c).

The spatial locations of V1-like RFs used to simulate preference-switching and preference-maintaining neurons (Fig. 7) were the same as previously detailed for center-surround RFs. *θ*_*c*_ (RF orientation angle) for each of the 53 units was chosen to match the orientation preferences reported in the imaging data (Fig. 7c, middle-left panel), instead of randomly chosen as in all other simulations of V1-like RFs.

### Stimuli (Fig. 1b)

#### All simulations

Stimuli were 2D static sinusoidal gratings presented in several aperture types with hard boundaries, with the gratings being defined as in Equations 2 and 3.

Twelve grating orientations (*θ*_*c*_ = 0,15,30,…,165^°^) and four grating phases (*ϕ*_*g*_= 0,90,180270 ^°^) were used to probe orientation selectivity. Multiple stimulus phases were chosen to minimize potential effects of phase-dependent RF responses. Gratings were presented at 100% contrast on a gray background. Minimum luminance was assigned a value of −1, maximum luminance was assigned a value of 1, and the gray background was assigned 0. The FOV of the entire stimulus was constructed to be 120° x 120° in visual field coordinates, corresponding to a sampling area of 200 × 200 pixels. Simulated stimuli never extended beyond this FOV.

#### Aperture simulation

Circular apertures had a radius of either 20, 30, or 40°. Square or diamond (square rotated 45°) apertures had a side length of 40°. Spatial frequency of the gratings ν_*g*_ was 0.04 cycles/° (Fig. 3a, top row).

#### Spatial frequency simulation

Circular apertures had a radius of 30°. Grating frequencies were chosen from six possible spatial frequencies (ν_*g*_ =0.01,0.02,0.04,0.16,0.32 cycles/^°^) (Fig. 4a, top row).

#### RF properties simulation

Circular apertures had a radius of 30°. Horizontal and vertical edge aperture conditions were made by masking either the bottom or left half of the circular aperture. Spatial frequency of the gratings ν_*g*_ was kept at 0.04 cycles/° (Fig. 3a, top row, center). For the investigation of preference-switching and preference-maintaining neurons (Fig. 7), 36 evenly spaced stimulus orientations were used.

### RF model responses

#### *RF responses in visual coordinates* (Fig. 1c)

The scalar response to each stimulus from each RF was calculated by performing a dot product between the vectorized images of each RF and stimulus. Responses were then normalized between 0 and 1 across all RF locations. This normalized response simulated the change in response from baseline driven by the presented visual stimuli. Independent Gaussian noise (mean: 0, standard deviation: 0.01) was then added to each normalized response. The total simulated response was calculated as follows:

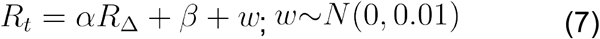

where *R*_*t*_ denotes the total response, *R* _Δ_ denotes the normalized, stimulus-driven change in response, *β* denotes a baseline response, *w* denotes the Gaussian distributed noise, and *α* denotes a scaling factor on the stimulus-driven change in response. Throughout this study, the baseline response *β* was always set to have a value of 1, and was also always set to have a value of 1. This resulted in a maximum possible increase (before adding noise) from baseline response of 100% (*α* /*β* =1). If a different maximum increase were desired, the ratio of *α* to *β* could be changed correspondingly (e.g., *α* /*β* =0.75 for a 75% maximum increase).

The matrix of responses for the set of center-surround RFs was a 4D matrix of size **12** (stimulus orientations) x **4** (stimulus phases) x {58,081, 103,041, 160,801} (RF locations for either ±30°, ±40°, or ±50° RF grids) x **2** (RF phases), while the matrix of V1-like RF responses had an additional 5^th^ dimension of size **8** for each RF orientation. For center-surround RFs, responses across RF phases were collapsed by thresholding negative responses to 0 and summing the On-center and Off-center responses together. For V1-like RFs, responses across RF phases were collapsed by squaring quadrature phase responses and summing the two together, which is the formalized method to achieve phase-invariant energy responses for Gabor RFs^37^. Collapsing responses across RF phases resulted in a matrix of size **12** x **4** x {58,081, 103,041, 160,801} for center-surround RFs or **12** x **4** x {58,081, 103,041, 160,801} x **8** for V1-like RFs. Responses were then averaged across stimulus phases, resulting in a matrix of size of either **12** x {58,081, 103,041, 160,801} for center-surround RFs or **12** x {58,081, 103,041, 160,801} x **8** for V1-like RFs.

#### *RF responses in anatomical coordinates* (Fig. 1d)

To compare model results to prior imaging data^9^, we developed a method for simulating model responses in anatomical coordinates. RF responses were approximated at anatomical locations using the previously simulated RF responses in visual coordinates. This process can be summarized in two steps: 1) defining a set of corresponding anatomical and visual coordinates, and 2) generating responses for those coordinates.

To define anatomical coordinates, we generated an evenly-spaced grid of coordinates in anatomical space. Natural neighbor interpolation^75^ was used to learn an approximate transform from anatomical to visual space coordinates (see following paragraph for brief overview). This was implemented by training two nonlinear interpolant functions that estimated x- or y-coordinates in visual space given pairs of x-y coordinates in anatomical space. Using prior retinotopic maps^9^ (Fig. 2a), we trained these two functions by designating 10 points in anatomical space as training features and the 10 corresponding points in visual space as training targets (Fig. 2b, white points). The extent of the square anatomical window (Fig. 2a, bottom) was defined as ±0.83 mm, and the extent of the square visual field grid (Fig. 2a, top) was defined as ±40°, following the scale illustrated in the retinotopy. We defined an evenly spaced grid (241 × 241 samples) of anatomical coordinates, *C*_*anat*_ (Fig. 2c, top, white points), overlaying the anatomical map. The two nonlinear interpolant functions were then used to find visual coordinates, *C*_*anat* → *vis*_ (Fig. 2c, bottom, white points), corresponding to *C*_*anat*_.

In natural neighbor interpolation, given a set of training features in the input space (e.g., a set of x- and y-coordinates in anatomical space) and a set of training targets in the output space (e.g., a set of x-coordinates or a set of y-coordinates in visual space), a nonlinear function is learned that transforms any feature in the input space to a corresponding target in the output space. As opposed to linear interpolation, the nonlinearity afforded by natural neighbor interpolation better approximated the nonlinear transform from anatomical to visual coordinates. The benefit of a more complex, nonlinear transform can be qualitatively appreciated when comparing anatomical to visual space from the prior imaging study^9^ (Fig. 2a, note distortion of the square grid in anatomical space).

To generate RF responses at coordinates *C*_*anat* → *vis*_, a linear weighting was performed on previously simulated RF responses (as described in *RF responses in visual coordinates*). The response at any coordinate *C*_*anat* → *vis*_ was estimated as a weighted average of RF responses from the 4 closest surrounding coordinates from the previously simulated grid of RF responses (e.g., Fig. 2c, bottom panel, RF responses at the 4 black points surrounding the top-leftmost white point). The weighted contribution from each surrounding RF response was determined by the spatial distance to the coordinate *C*_*anat* → *vis*_. Any *C*_*anat* → *vis*_ coordinates falling outside the range of the previous RF response grid were excluded. For V1-like RFs, weighted responses at coordinates *C*_*anat* → *vis*_ were first calculated for each possible RF orientation (of 8), and afterwards, the response from a random RF orientation was chosen at each coordinate location.

#### *Population RF responses* (Fig. 1e)

We used spatial smoothing to approximate population responses. A 2D Gaussian smoothing kernel was applied across responses simulated at anatomical coordinate locations *C*_*anat*_, with a standard deviation of 0.0083mm, or 0.44° in visual coordinates (on average, owing to a nonlinear transform between anatomical and visual space). In terms of RF coverage, responses from a grid of approximately 16-25 RFs (4×4 to 5×5) would fall within 1 standard deviation of the smoothing kernel.

#### *Single-unit orientation tuning* (Fig. 7e)

Single-unit tuning functions were obtained by simulating responses of either a single center-surround or V1-like RF to 36 evenly spaced orientations from 0-180° and fitting the set of responses with a circular von Mises function, using the Circular Statistics Toolbox through Matlab^76^. The curve was duplicated from 180-360° to visually match the tuning curves obtained from empirical direction selectivity measurements (Fig. 7d).

### Orientation preference maps and orientation selectivity index

Global orientation selectivity indices (gOSI) were calculated based on the following equation,

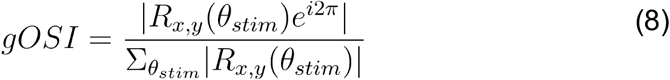

where *θ*_*stim*_ denotes stimulus orientation, and *R* denotes the matrix of responses across RF locations. We normalized gOSI from 0-1 across the set of responses from each RF across all stimulus orientations for each stimulus manipulation (i.e., size, shape, spatial frequency). We used gOSI as a transparency and thresholding value in our orientation preference maps to highlight RFs based on their degree of orientation selectivity.

## Acknowledgements

This research was supported by the Intramural Research Program of the NIMH (ZIAMH002966) to E.P.M. and the Wu Tsai Neurosciences Institute at Stanford to J.L.G. Support to A.K. is provided by the Graduate Partnerships Program at the NIH in conjunction with the Stanford Neurosciences Interdepartmental Program.

## Author contributions

A.K.: conceptualization, methodology, investigation, validation, formal analysis, visualization, writing – original draft, writing – review and editing

J.L.G.: conceptualization, methodology, investigation, validation, resources, supervision, project administration, funding acquisition, writing – review and editing

E.P.M: conceptualization, methodology, investigation, validation, resources, supervision, project administration, funding acquisition, writing – review and editing

## Competing interests

The authors declare no competing interests.

## Notes

### Competing Interest Statement

The authors have declared no competing interest.

### Summary of Updates

Abstract and introduction section updated to mirror latest changes for submission to peer-review

